# Life history strategies and niches of soil bacteria emerge from interacting thermodynamic, biophysical, and metabolic traits

**DOI:** 10.1101/2022.06.29.498137

**Authors:** Gianna L. Marschmann, Jinyun Tang, Kateryna Zhalnina, Ulas Karaoz, Heejung Cho, Beatrice Le, Jennifer Pett-Ridge, Eoin L. Brodie

## Abstract

Efficient biochemical transformation of belowground carbon by microorganisms plays a critical role in determining the long-term fate of soil carbon. As plants assimilate carbon from the atmosphere, up to 50% is exuded into the area surrounding growing roots, where it may be transformed into microbial biomass and subsequently stabilized through mineral associations. However, due to a hierarchy of interacting microbial traits, it remains elusive how emergent life-history strategies of microorganisms influence the processing of root exudate carbon. Here, by combining theory-based predictions of substrate uptake kinetics for soil bacteria and a new genome-informed trait-based dynamic energy budget model, we predicted life history traits and trade-offs of a broad range of soil bacteria growing on 82 root exudate metabolites. The model captured resource-dependent trade-offs between growth rate (power) and growth efficiency (yield) that are fundamental to microbial fitness in communities. During early phases of plant development, growth rates of bacteria were largely constrained by maximum growth potential, highlighting the predictive power of genomic traits during nutrient-replete soil conditions. In contrast, selection for efficiency was important later in the plant growing season, where the model successfully predicted microbial substrate preferences for aromatic organic acids and plant hormones. The predicted carbon-use efficiencies for growth on organics acids were much higher than typical values observed in soil. These predictions provide mechanistic underpinning for the apparent efficiency of the microbial route to mineral stabilization in the rhizosphere and add an additional layer of complexity to rhizosphere microbial community assembly.

## Introduction

Trait-based approaches to reconcile the complexity of microbiomes triangulate conceptual theory, analytical measurements, and numerical models in order to provide mechanistic explanations of ecological patterns and robust predictions of ecological dynamics and function [28]. High-throughput genome sequencing and concurrent advances that help infer functional traits from genomes have increased the availability of large-scale microbial functional trait datasets from diverse habitats [55]. However, advancing our understanding of functional traits from genomes alone is challenging, because each organism’s traits interact within an ecological context in response to a changing environment (e.g., substrate availability) and the presence of other organisms (e.g., through competition and mutualism) [38]. Unlike the expansion of genomic information, compilations of microbial phenotypic trait data have tended to accumulate at a slower pace [34]. As most soil bacterial species remain uncultured, the vast majority of soil bacteria remain known only from their genomes [61]. For this reason, theoretical approaches are needed to enable the integration of microbial functional traits with the dynamics of a soil ecosystem in order to predict microbial phenotypes and their consequences for soil biogeochemistry. In this regard, dynamic systems models of energy and mass exchange between organisms and their environments [24] offer a promising and tractable procedure for data-model integration that moves beyond genetic potential and allows us to characterize the realized phenotypes of microbes.

Trait integration is inherently hierarchical and scale-dependent: genes integrate into metabolic pathways, whose interactions determine phenotypes, which combine into broad emergent properties that define life history strategies, and finally, these strategies dictate population dynamics through environmental feedback and community composition [20]. At the same time, it is important to recognize that many traits are highly integrated because of their intrinsic dependence and synergy with other traits [1]. Some microbial traits vary with taxonomy due to evolutionary influences. In those cases, differences among species and taxonomic groups (e.g., family, order) can explain a large amount of variation in trait values [40]. Evolutionary history thus enables microbial ecologists to predict some community-level mean trait values and ecosystem functions across environmental variation. However this is not universally true, and other traits, for example genomic traits related to the capacity to use simple carbon substrates, are highly phylogenetically dispersed. Since common phylogenetic biomakers (such as the 16S rRNA gene) can only resolve up to 50 million years of evolution [44], recently acquired traits (such as those acquired through horizontal gene transfer) may not be not predictable by phylogeny, yet may be key to local niche adaptation [6]. This weakens associations between phylogeny and function, and complicates our ability to incorporate microbial biodiversity into models.

To facilitate the assignment of functional attributes to microbial communities, recent studies have classified individual genomes based on quantitative biophysical and life history traits, as well as biochemical similarities [68]. These results challenge binary classifications of microbial life-history strategies -that are rooted in r-K selection theory [70], and question the use of mean trait values to describe groups of genomes with shared metabolic capabilities [50]. Biophysical traits and life history traits (e.g., genome size, minimum generation time) can vary widely within substrate-use pathways [69]. Indeed, a major portion of fitness at intermediate levels of trait integration is determined by how all the traits of a particular body plan interact within fundamental biophysical constraints [26]. In accordance with the metabolic theory of ecology, cell size is a highly integrated phenotypic trait that links the principal axes of metabolic rates, resource acquisition, and stress tolerance [11]. The idea that metabolic rate determines population processes and life history is well accepted in other branches of ecology [12]. However for microbes, there is no obvious connection between the scaling of any single cellular feature and overall metabolic rate [27]. Nonetheless, metabolic rate explicitly constrains trade-offs between cellular functions during exponential growth [25]. Accounting for cellular trade-offs based on the combined variance in biophysical and life history traits may therefore allow identification of microorganism niche preference based on interacting life history strategies, and substrate preferences.

Here, we present a genome-informed, trait-based dynamic energy budget model (DEBmicroTrait) to integrate genome predicted traits and their interactions within a dynamic environment, letting life history strategies and niches of soil bacteria emerge from fundamental thermodynamic, biophysical, and metabolic principles that constrain trait variation, trait linkages and ultimately organism fitness (Suppl. Info. A-D). We have focused on integrating genomic traits that distinguish soil bacteria that actively consume plant-derived carbon in the rhizosphere (the area surrounding growing plant roots) relative to other bacteria that may have a fitness advantage in neighboring bulk soils [73]. The rhizosphere is chemically diverse and a critical hotspot for biogeochemical transformation with high potential for carbon stabilization through microbial carbon assimilation and subsequent mineral-surface stabilization [45, 58]. To better understand interactions between life history traits and substrate-use pathways in the rhizosphere, we simulated the growth of 39 bacterial isolates on 82 plant exudate metabolites. We benchmarked simulations quantitatively and qualitatively against (i) genome predicted minimum generation times that agree well with measured growth rates of isolates, and (ii) substrate uptake preferences of isolates given as a percentage of metabolite depletion in a mixed growth medium [73]. We then simulated population-level estimates of key traits, including realized growth rate and carbon assimilation rate, across different metabolite classes that are known to be rhizosphere exudates. By enabling integration across multiple levels of trait hierarchy, we represent trait relationships dynamically, rather than relying exclusively on fixed categories or prescribed trait correlations. In this context we can then assess the distribution of predictive power across the hierarchy of genome-inferred traits, on microbial phenotypes. Our hierarchical trait integration allows us to observe emergent strategies for energy and resource acquisition and allocation in microbial metabolism, and explain these in terms of trade-offs between growth rate (power) and growth efficiency (yield) that ultimately determine microbial fitness in communities [52]. These interactions -among multivariate trait strategies-can then be mapped to generalizable rules for how microbes interact with plant traits, acquire complex resources, and regulate soil organic matter formation.

## Results

Growing roots dramatically alter the chemical and physical habitat for microorganisms and exude photosynthesis-derived carbon that can be broadly classified into sugars, amino acids, organic acids, fatty acids, nucleotides and auxins [5]. In fact, root exudation itself might be a generalizable trait with rates and composition that can be predicted from plant functional traits [72], with consistent temporal patterns across plant development stages [73]. The chemical composition of root exudates interacts with microbial metabolite preferences that are predictable from genomic traits (Figure 1). These soil microorganisms differ in genomic traits related to resource acquisition, namely their metabolic potential to utilize organic acids (Figure 1B) as well as their potential for plant polymer degradation by glycoside hydrolase enzymes (Figure 1C). Contrary to expectations [16] and conventional wisdom, previously we found that many of the bacteria that responded positively to root growth had slower maximum specific growth rates (both in laboratory growth experiments, as well as minimum generation times predicted based on genomic signatures (Figure 1A). We hypothesized that substrate preference along with substrate utilization efficiency interact to confer a growth efficiency-based fitness advantage for many bacteria in the rhizosphere. However, growth efficiency can vary widely with limiting resource concentration and the free energy content of chemical compounds that are released by plants [52]. These patterns are overlaid with physiological variations in resource use between bacteria [41, 48, 54] that likely affect phenotypic trade-offs such as carbon-use efficiency (CUE) and combine to influence the timing of rhizosphere community succession.

**Figure 1.**
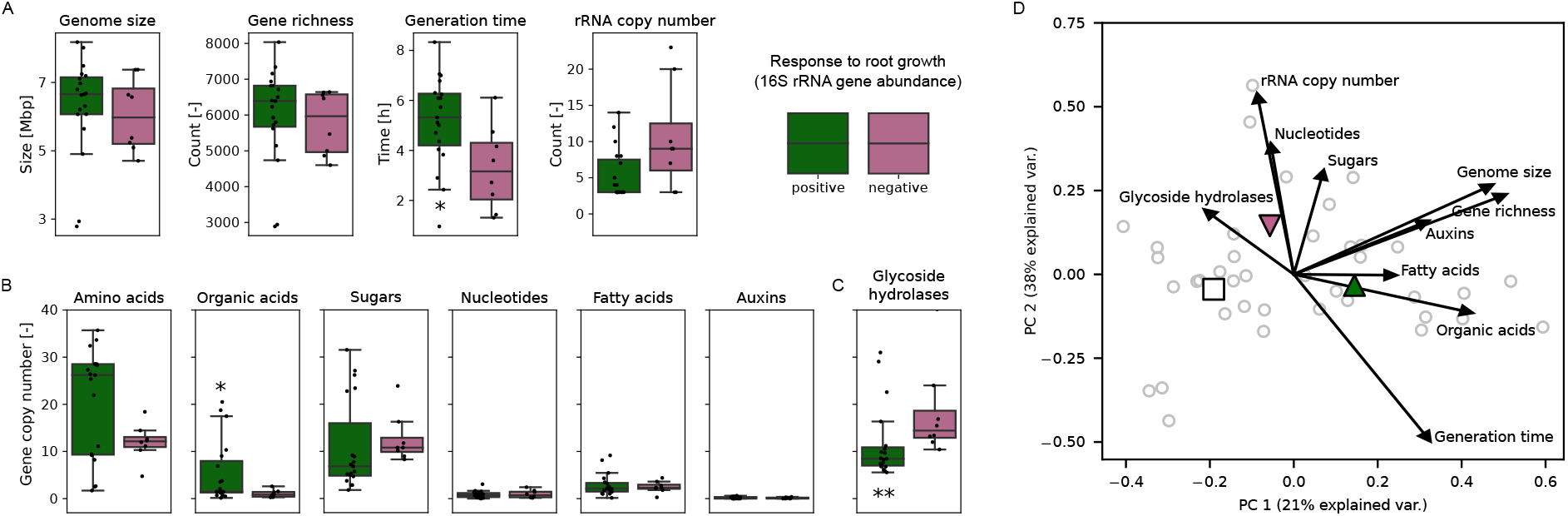
Genomic trait distributions of soil bacterial isolates classified based on the response to root growth of *Avena barbata*. Modified from [73]. **A** Genome size, gene richness, minimum generation time, and rRNA operon copy number predicted from genome sequences of soil isolates. **B** Monomer transporters. **C** Glycoside hydrolase enzymes. Gene copy numbers for monomer transporters and glycoside hydrolases were normalized by genome size. Differences in genomic trait distributions between positive and negative responders were evaluated using the Kruskal-Wallis one-way analysis of variance, and traits with significant differences are annotated according to the R project standard convention [49] used throughout the manuscript (*** p<=0.001, ** p<=0.01, * p<=0.05). Isolate response groups were classified based on changes in 16S rRNA gene abundance over the plant developmental stages of *Avena barbata*. In each boxplot, a point denotes a single isolate. The top and bottom of each box represent the 25th and 75th percentiles, the horizontal line inside each box represents the median and the whiskers represent the range of the points excluding outliers. Positive responders: n=19, negative responders: n=8. **D** Principal component analysis illustrating covariations among genomic traits shown in panels (A)-(C). Highlighted symbols represent average coordinates of positive, negative, and undefined (n=12) isolate response groups.

### Substrate Preference in the Rhizosphere

Genomic traits provide information on the constraints and demands for substrate uptake of soil isolates. We synthesized these by considering interactions between cell size and cellular carbon density, cell surface area-to-volume ratio and growth rate potential in the equilibrium chemistry approximation (ECA) for nutrient uptake [63]. Similarly to Michaelis-Menten kinetics, ECA kinetics are defined by a maximal reaction rate (*V*_*max*_) and an apparent half-saturation constant (*K*). Both *V*_*max*_ and *K* can vary widely with cell size and accessible substrate binding sites on the cell surface [64], permitting us to explore the variation and allometric scaling of both kinetic parameters by reference to the transport potential required to support a given genome-inferred maximum specific growth rate [19]. The estimated binding-site densities were benchmarked against a compilation of existing data on nutrient transport and then scaled by relative gene frequencies of transporter genes in order to account for the evolutionary history of substrate preference encoded in rhizosphere genomes (Suppl. Info. B).

Accordingly, we find that at high substrate concentrations, the substrate preference for organic acids of positive rhizosphere responders (Figure 1B) is reflected in modeled maximum specific uptake rates (Figure 2A). Negative responders, achieve significantly higher maximum specific uptake rates to match genome-predicted maximum specific growth rates (Figure 1A), except for organic acids and auxins. Maximum specific uptake rates exhibit an emergent trade-off against substrate half-saturation constants of soil isolates. With half-saturation constants in the order of typical plant metabolite concentrations (Figure 2B), positive responders can respond quickly to small temporal changes in exudation rates [46]. Independent of metabolite class, the overall shape of the trade-off is concave (Suppl. Figure S3 A) and also reflects uptake phenotypes that are not specialized to either fast uptake (high *V*_*max*_) or rapid response (low *K*). These uptake generalists are likely competitive across a wider range of substrate concentrations in the rhizosphere [35].

**Figure 2.**
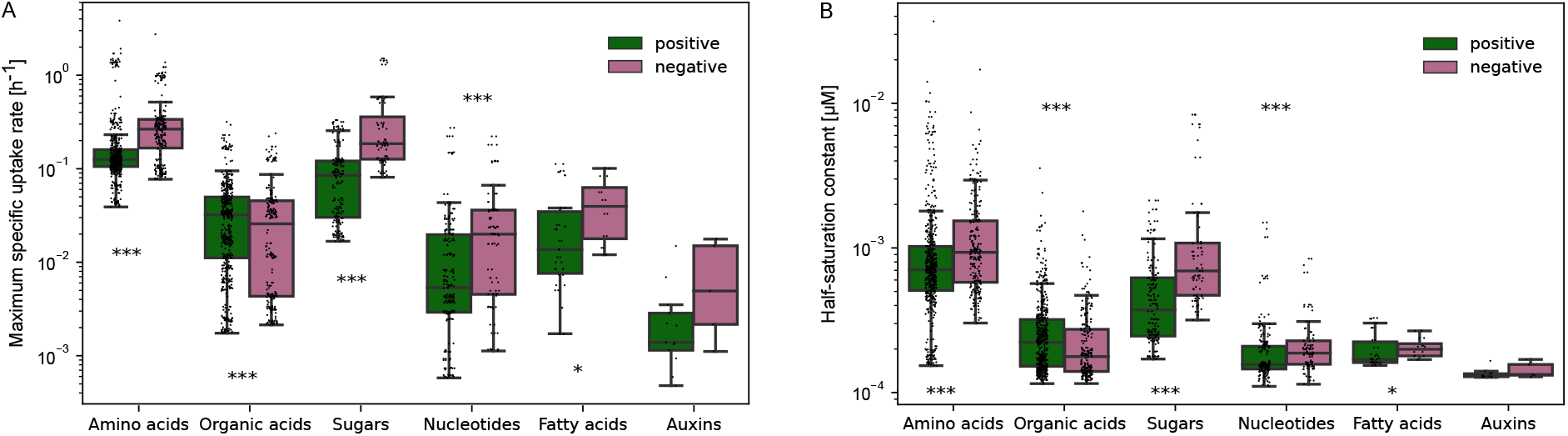
Theory predicted distributions of root exudate metabolite uptake parameters of soil bacterial isolates. **A** Maximum specific uptake rate. **B** Half-saturation constant. Differences in uptake trait distributions between positive and negative responders were evaluated using the Kruskal-Wallis one-way analysis of variance. In each boxplot, a point denotes a single substrate-consumer relation in the equilibrium chemistry (EC) approximation [64, 65]. The top and bottom of each box represent the 25th and 75th percentiles, the horizontal line inside each box represents the median and the whiskers represent the range of the points excluding outliers. Substrates: n=82, consumers: n=27.

### Phenotypic Comparisons and Trade-offs

Resource acquisition traits are only one facet of ecological strategy variation. We thus further analyzed biomass production (BP) and respiration (BR) rates of isolates in order to explore relationships between realized growth rate and CUE, calculated as BP/(BP+BR), across 3198 batch simulation runs. Simulated cellular rates, spanning nanomolar ranges at target cell densities between 10^5^ and 10^6^ cells ml^-1^, are on the high end of commonly observed values for bacteria [41]. Corresponding values for the emergent CUE range from 0.07 to 0.74, with a median of 0.49, suggesting that on average approximately half of the consumed carbon is typically lost via respiration (Suppl. Figure S6 A). The average simulated CUE is close to the average observed value of 0.55 in soil [37], which itself is not far from the average maximum value of 0.6 in pure culture studies [57]. We then partitioned variation in CUE based on isolate identity and metabolite type (Suppl. Table S1). Within taxonomy, narrower taxonomic groups (e.g., class, species) explained more of the variation than broader levels (e.g., phylum). Specifically, as much as 20% of the variation in CUE across metabolites was conserved at the class level, and an additional 18% could be explained by species identity. Yet, metabolite type was a strong predictor of CUE across species (48%), and explained 33% more variation than differences in the mean Gibbs free energy and stoichiometry based on metabolite class (15%). Within a given metabolite class, more than half of the variation in CUE (54%) was due to species identity. Although some metabolite classes had significantly different median values for CUE (Suppl. Figure S8), the standard deviation within each metabolite class ranged from 0.09 to 0.15 and was therefore similar to the standard deviation across class medians (0.09). Feature contributions to CUE were calculated using isolates classified previously as undefined rhizosphere responders, as out-of-bag samples, in a random forest regression algorithm [62]. Based on mean decrease in the model prediction accuracy (Suppl. Figure S7), traits related to carbon assimilation had the largest influence on prediction accuracy of CUE (58%), reflecting energetic and stoichiometric constraints that depend on both metabolite- and isolate-specific characteristics. Differences in protein synthesis efficiency (19%) and extracellular enzyme production (4%) explained a large amount of inter-species variation in CUE. Taken together, CUE emerges as a complex physiological trait that can be modified by metabolite chemistry, resulting in significantly higher CUE for positive responders for growth on amino acids, organic acids, sugars, and auxins (*p <* 1*e*^−3^, Suppl. Figure S8).

We then tested for trade-offs between growth rate and CUE (Figure 3A). For metabolites that are assimilated at high rate and yield (sugars, amino acids), a significant inverse relationship exists between growth rate and CUE (F_1,3_=13.83, r^2^ =0.82, p=0.03). The model predicts a 7.6% decrease in CUE for every per minute increase in growth rate. Maximum CUE is achieved for positive responders growing on glucose, while maximum growth rates occur at sub-optimal CUE. Inefficiencies of the microbial growth machinery at high growth rates are explained in part by accelerated dilution of storage compounds due to (volume) growth (Suppl. Info. Eq. 3, [15]), as well as decreasing structural biomass yield with increasing protein synthesis rates (Suppl. Figure S4 C, [39]). At comparatively lower growth rates on the other hand, for growth on organic acids, fatty acids, nucleotides and auxins, we found a positive relationship between growth rate and CUE (F_1,11_=7.74, r^2^=0.41, p=0.02). Here, the model predicts a 13% increase in CUE with every minute increase in realized growth rate. Differences in the slope of the scaling relationship between growth rate and CUE in the two growth regimes can be understood by analyzing correlations between BP and BR in isolate metabolism (Figure 3B). In the low growth rate regime, BP and BR scale isometrically (with slope *β* ∈ [0.95, 1.05], t_1332_=0.06, p=0.95), indicating that respiration is a major control on biomass production and CUE. Indeed, maintenance requirements, including extracellular enzyme production, explain most of the variability in the growth strategies of isolates at low growth rates (Figure 3D). At high growth rates on the other hand, BP and BR scale allometrically (with slope that is significantly different from one, *β* ∈ [0.56, 0.63], t_1437_=-24, p<1e-5). Energy budget allocation to maintenance, extracellular enzyme production and assimilation are nearly independent (orthogonal) axes of variation, while growth production correlates with higher biomass turnover (Figure 3C). A Kruskal-Wallis test showed that the principal components of the different response groups are significantly different (H_PC1_(2)=329, H_PC2_(2)=105, p<2.2e-16). A post-hoc test using Dunn’s test with Benjamini-Hochberg correction confirmed the significant differences between positive and negative responders along PC1 (PC2) (p<2e-16 (<9.3e-4)) and negative and undefined responders (p<1.6e-5 (<2e-16)) in the high growth regime. In the low growth regime, the ecological strategies of negative and undefined responders coalesce along both principal components, p=0.71 (0.054). They differ significantly from positive responders that showed overall higher maintenance rates due to larger cell sizes, but allocate a smaller fraction of the total energy budget to cellular maintenance processes (p<2.5e-5, Suppl. Figure S9 A).

**Figure 3.**
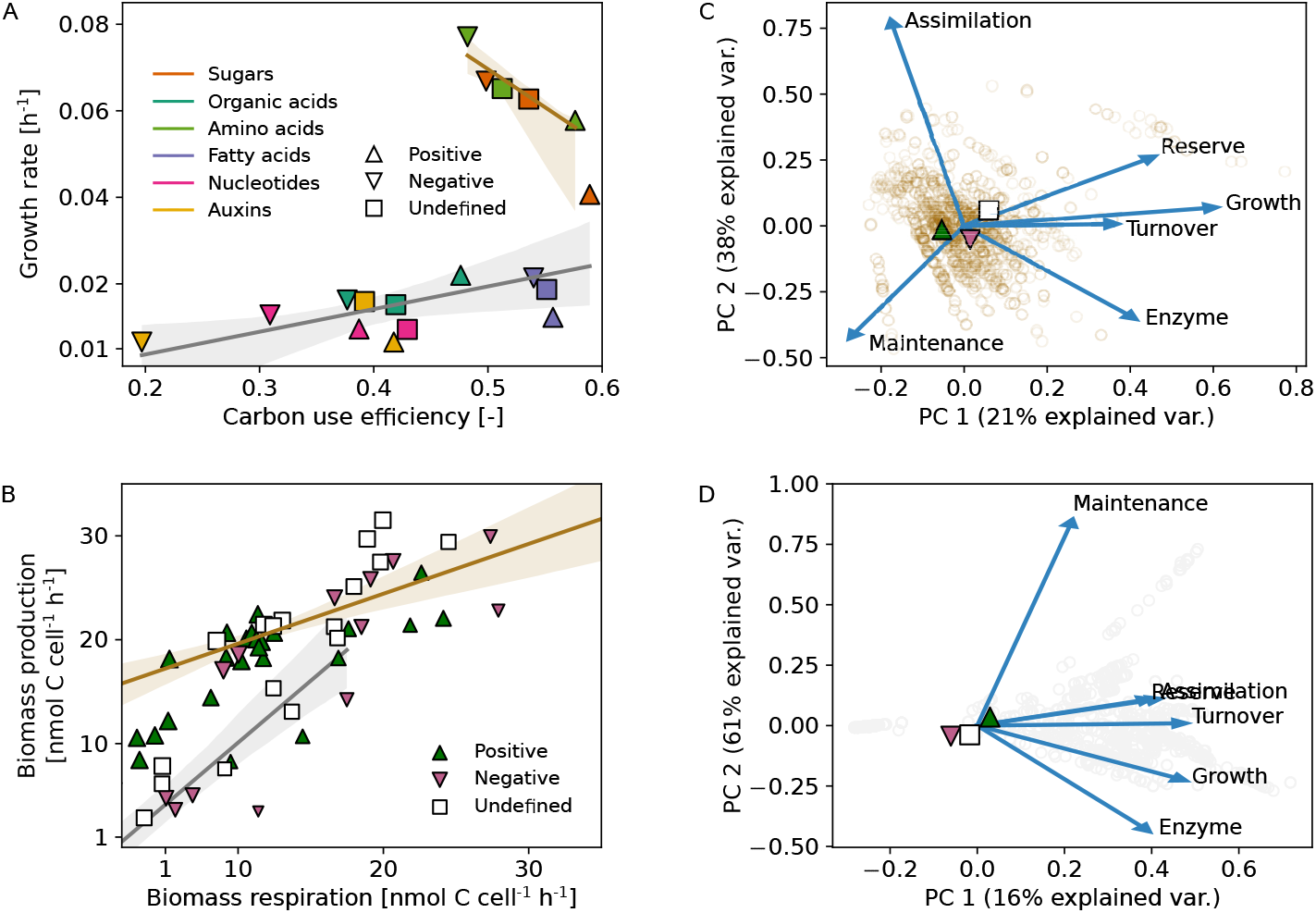
Phenotypic traits and trade-offs during batch growth on root exudate metabolites. **A** Relationships between realized growth rate and carbon-use efficiency. Median trait values are plotted using different colors and shape depending on metabolite class (sugars, organic acids, amino acids, fatty acids, nucleotides, auxins) and isolate response to plant root growth (positive, negative, undefined). **B** Relationships between biomass production and respiration rates. Symbol size is scaled by carbon-use efficiency. Solid lines indicate the observed regression lines, while shaded areas indicate the 95% confidence bands. Throughout plots **A-D**, two distinct growth regimes are distinguished by brown (high growth regime) and gray (low growth regime) colors. **C** Principal component analysis illustrating covariations among modeled fluxes delineating growth strategies of isolates at high and **D** low realized growth rates. Highlighted symbols represent average coordinates of positive, negative, and undefined isolate response groups. Substrates: n=82, consumers: n=39, simulations: n=3198.

### Relationships with Genomic Traits

Model predicted growth rates of the bacteria span two orders of magnitude across plant root metabolites, ranging from 0.0044 h^-1^ to 0.46 h^-1^ (Suppl. Figure S6 B). Growth rates at the lower end of the low growth rate regime (0.0044 - 0.039 h^-1^) are reflective of growth rates of autochthonous bacteria occurring in unamended soils as previously determined by quantitative stable isotope probing using ^18^O-labeled water [30], while growth rates in the high growth rate regime span typical values observed in pure cultures under laboratory conditions [53, 66]. Predicted growth rates were partially confirmed with measured growth rates at carbon concentrations representative of the original growth medium used to inoculate the bacterial isolates ([73], Suppl. Figure S10). The largest model deviations (r^2^=0.67, RMSE = 0.1 h^-1^) occur at the lower end of observed growth rates, where the model underpredicts the observed values. Interestingly, very few isolates reach their maximum growth potential for simulated growth on root exudate concentrations (125 mg C per liter) corresponding to concentrations of dissolved organic carbon detected in soil from the University of California Hopland Research and Extension Center (Hopland, CA, USA; 38°59^*′*^ 34.5768^*′′*^ N, 123°4^*′*^ 3.7704^*′′*^ W), where these bacteria were originally isolated (Suppl. Figure S11). Nonetheless, realized growth rates and CUE are strongly correlated with the number of copies of the 16S rRNA gene at high growth rates (Suppl. Table S2), where this single genomic trait explains about a third of the variation (r^2^=0.30 (0.44)). Isolates with more copies of the rRNA gene grow faster on glucose and amino acids, 0.0125 h^-1^ per additional gene copy. Here, CUE is inversely related to growth rate and decreases by 0.015 units per rRNA copy number. Relationships with genome size are only significant in interaction with rRNA copy number, where genome size has been shown to be positively correlated with the number of rRNA gene copies [53]. At low growth rates on the other hand, more than 96% of the variance in growth rates is unexplained by rRNA copy number and genome size. It thus appears that at lower growth rates, substrate limitation occurs from environmental supply, whereas at high growth rates, substrate limitation occurs inside the cell as caused by growth-induced dilution [21]. However, both genomic traits are relatively good predictors of CUE (r^2^=0.35), where CUE declines by 0.02 (0.01) units per rRNA gene copy number (Mbp in genome size).

### Interactions between Substrate Preference and Carbon-Use Efficiency

Next, to determine whether substrate preferences of isolates interact with CUE to confer a selective advantage in the rhizosphere, we analyzed mixed media simulations for differences in root exudate metabolite uptake across positive and negative responders to root growth. We found that 39 out of the 82 simulated exudate metabolites show non-negligible differences in metabolite uptake between positive and negative responders (Figure 4). For 16 out of the 39 metabolites that had previously been identified experimentally [73], we found the largest cumulative differences in the uptake of positive responders for plant hormones (indole-3 acetic acid, abscisic acid), followed by a cluster of aromatic organic acids (caffeic, shikimic, 3-dehydroshikimic, trans-cinnamic, salicylic, nicotinic). Nucleosides, on the other hand, are more preferentially consumed by the negative responders. Differences in uptake of 16 out of the 39 metabolites agree qualitatively with uptake of the specific compounds from the growth medium measured by LC-MS [73]. We then tested for differences in CUE for growth on metabolites that are preferentially consumed by either response group (Figure 4 inset). For metabolites that are preferentially consumed by positive responders, there are significant differences in CUE between positive and negative responders (H(2)=23, p<1e-5). Metabolites that are preferentially taken up by negative responders show a wider range in CUE, with median values that are indistinguishable between the bacterial response groups, but significantly lower than for growth on metabolites that are preferentially consumed by positive responders (H(2)=6, p=0.01).

**Figure 4.**
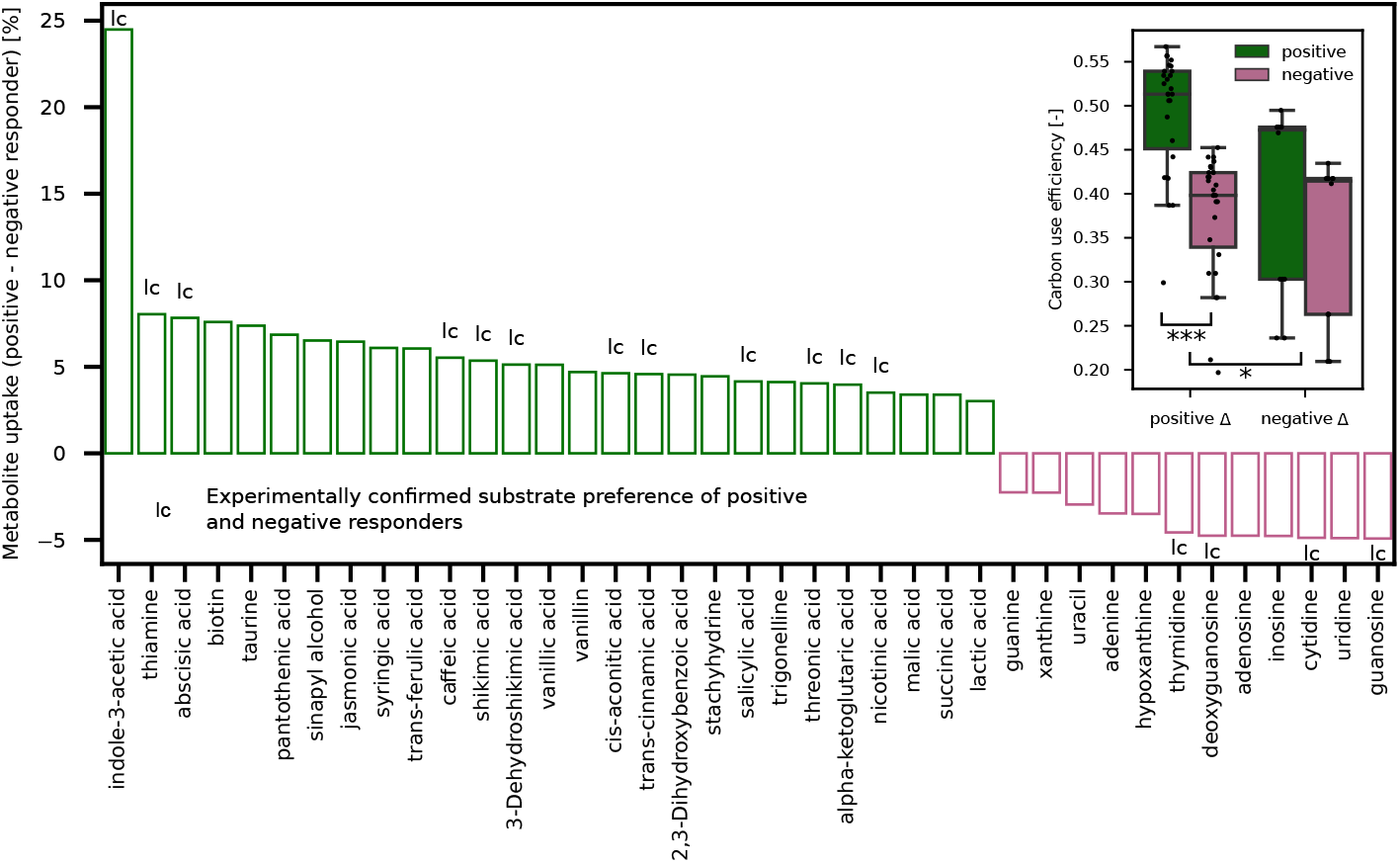
Substrate preference and carbon-use efficiency in mixed root exudate metabolite medium. Metabolites with the largest differences in uptake are shown on the x-axis (n=39). Each bar corresponds to differences in metabolite uptake quantified as a percentage of the difference in metabolite depletion from the medium. Metabolite uptake preferences that were confirmed experimentally [73] are denoted by the letters ‘lc’. The figure inset shows differences in carbon-use efficiency for substrates that were preferentially consumed by positive responders (positive Δ, n=27) and negative responders (negative Δ, n=12). Differences in carbon-use efficiency within and between metabolite preference group were evaluated using the Kruskal-Wallis one-way analysis of variance. In each boxplot, a point denotes a single metabolite-isolate pair. The top and bottom of each box represent the 25th and 75th percentiles, the horizontal line inside each box represents the median and the whiskers represent the range of the points excluding outliers. Substrates: n=82, consumers: n=27, simulations: n=27.

## Discussion

### Power-yield Signatures of Rhizosphere Succession

Many specific trade-offs have been demonstrated, in individual microorganisms or populations, that can be described as informed choices between power and yield [3, 8,17]. Recently, an extensive analysis of microbial growth thermodynamics revealed that growth rate and yield do not show such a trade-off under energy-limited conditions [14], and there exists insufficient evidence that fast growth universally reduces growth efficiency, both in pure culture studies [41, 48, 52] and whole-soil communities [32, 74]. Conceptual theory suggests that realized growth rate is in fact not a coherent ecological strategy, but rather a complex emergent property that depends on both growth yield and the rate of resource acquisition [36]. Across time, it has been observed that microbial communities adjust power-yield trade-offs depending on the constraints on the system [51]. Maximum power is significant early in colonization events, when rapid growth is called for and when the environment provides an abundance of free energy from resources [32]. Conversely, when resource availability is low, biological assemblages will form that decrease energy dissipation resulting in proportionally higher growth yields [23].

Along the same lines, our results show that growth rate and yield diverge even within yield-optimized strategies, as high yield can be achieved with different growth rates (Figure 3A). Across the plant growing season, soil microbial communities in the rhizosphere undergo succession to optimize the trade-off between power and yield in a predictable manner. Bacterial growth rate and CUE trade off during growth on metabolites exuded early during plant growth (sugars, amino acids) highlighting an early successional growth strategy where power is optimized over yield. The corresponding realized growth rates are strongly correlated with the number of rRNA copies in the genome (Suppl. Table S2), indicating that early successional growth rates of bacteria under nutrient-replete conditions are to a large part constrained by maximum growth potential [30]. The high growth rates are consistent with patterns of amino acid uptake by bacteria, where rapid uptake and subsequent incorporation into cell wall material result in short residence times of these compounds in soil environments [10]. Additionally, fast growth is strongly correlated with quick biomass turnover (Figure 3C) fueling the soil microbial carbon pump early during plant development [31].

Across time, we find that yield and resource acquisition strategies are tightly linked along gradients of resource availability, as bacteria grow more slowly on root exudates that are released during later plant developmental stages (organic acids, fatty acids, auxins). At these suboptimal growth rates, biomass production and respiration rates scale isometrically (Figure 3B) suggesting that respiration is a major control on biomass production and CUE [41]. The non-zero intercept of this relationship indicates that the fraction of energy budget allocated to maintenance of bacteria explains a large proportion of fitness at low growth rates. Additionally, density-dependent population dynamics such as microbial mortality [22], become less pronounced over time, as bacterial growth rate and biomass turnover are decoupled. Significantly lower biomass turnover in the low growth rate regime (Suppl. Figure S9 B) leads to a relationship between CUE and growth rate that is now positive. Additionally, the accumulation of energy storage compounds during early plant development (Figure 3C) may serve to help organisms produce biomass through efficient storage compound biosynthesis [71]. Together with CUE values that now span almost the whole range of values typically observed in soil [37], our results suggest that selection for efficiency is a primary driver of rhizosphere community composition late in the plant growing season.

### Carbon Stabilization through Resource Specialization in the Rhizosphere

Trait-based frameworks have successfully been used to confront ecological theory on resource-dependent succession in the host microbiome of a carnivorous pitcher plant [9]. Our results suggest qualitatively similar successional patterns across resource gradients, shifting over time from abundant resource acquisition to higher yield strategists, while also highlighting the complexity of substrate utilization in the rhizosphere. It appears that the ability to preferentially consume and grow efficiently on a handful of key plant root metabolites is a distinguishing feature of rhizosphere bacteria (Figure 4).

Based on our results, we posit that these ecophysiological controls interact with shifts in microbial density to influence pathways of soil organic matter formation across the plant growing season. As microbial density changes over the growing season, the release of sugars and amino acids, with a negative correlation between CUE and chemical polarity (a proxy for mineral sorptive affinity), coincides with rapid growth (more carbon overall into microbes), however the subsequent emergence of more yield optimized guilds (more carbon per specific microbe) may selectively enhance the stabilization of compounds through the deposition of senesced microbial biomass, that would otherwise be adsorped to mineral surfaces [59]. In particular, we find that, after the initial rhizosphere colonization by bacteria with power-optimized growth-strategies, it is the chemical succession of root exudate metabolites that selects for specific organisms with traits that result in higher CUE in the rhizosphere. Bacteria that responded positively to root growth had a 39% higher growth efficiency on average for organic acids with aromatic rings than those isolates that responded negatively to growing roots. Salicylic acid, a phenolic acid with high sorption potential to mineral surfaces, has been shown to correlate with specific taxa that are frequently enriched in the rhizosphere [4]. Likewise, we observed large differences in substrate preferences and CUE among isolates for plant hormones (indole-3-acetic acid, abscisic acid). Indole-3-acetic acid is an important signalling molecule in plant-bacterial interactions promoting plant growth [29] and the corresponding biosynthesis genes are frequently enriched in genomes of high-yield strategists that actively consume photosynthate in the rhizosphere [16]. Furthermore, CUE not only differed between isolate response groups in the rhizosphere, but our predicted growth efficiencies for these positive responders were also much higher than typical CUE values observed experimentally when phenolic or polyvalent organic acids, including salicylic acid, were added to bulk soils (i.e. 0%-30%, [59]). Taken together, these results strengthen prior evidence for direct manipulation of the soil microbiome through the specific composition of exudates [73] and provide mechanistic explanations for the apparent efficiency of the microbial route to mineral stabilization in the rhizosphere [58]. However, the mechanistic roadmap for how plant root carbon enters microbial and mineral-associated soil pools is even more complex. In particular, for senescing roots that have ceased exudation, microbial investment in extracellular polymer breakdown will further increase resource-based niche partitioning [42]. Further, positive interactions between rhizosphere bacteria may contribute to a more efficient microbial stabilization route via cross-feeding of metabolic byproducts [67], although the stability of microbially-derived carbon is likely impacted by the mechanism and kinetics of microbial mortality [60].

### Generalizable Trait Interactions in the Rhizosphere

While the rhizosphere bacteria considered here (n=39) cannot capture the full range of bacterial genomic or ecophysiological diversity, our modeling study fits within a long line of evidence demonstrating generalizable differences in life-history strategies of soil bacteria [2, 36]. In particular, we find that life history classifications rooted in r-K selection theory [18, 47] fail to fully describe ecological strategies in the rhizosphere, as various trade-offs between high yield, resource acquisition, and select stress tolerance traits (energy storage and maintenance) can, e.g., manifest in negative correlations between growth rate and yield during early plant development (Figure 3A).

Previous work using genome-scale metabolic modeling has predicted phylogenetic structuring of CUE at the subphylum level for over 200 environmental taxa [54]. However, empirical studies to validate phylogenetic signals of efficiency are limited as the number of growth substrates and organisms are typically insufficient for building a predictive model of CUE in unsampled regions of the phylogenetic tree [48]. The poor ability to predict CUE based on phylogeny, contrasts with experiments that report substantial variation in CUE among environmental isolates [41], commensurate with the amount of variation (13%-20%) that was attributable to differences among broad taxonomic groups considered here (Suppl. Table S1). In contrast to our findings, resource type was identified as a relatively weak predictor across species [41], suggesting that differences in resource supply alone will not affect CUE as much as changes that arise due to subsequent shifts in community composition and their use of those resources. Here, we find that detectable differences in CUE do exist in the averages between metabolite classes in the rhizosphere (Suppl. Figure S8). These differences are not outweighed by variation within classes, as standard deviations within metabolite classes are similar to those between classes. While mean CUE values for metabolite class in the rhizosphere reflect broad differences in the Gibbs free energy content and stoichiometry of metabolites released by roots, the quantitative trait variation within metabolite class conveys important information about substrate preference and competitiveness of individual organisms at different stages of the growing season.

Bacteria with large genomes have been shown to have a tendency to occupy a broader range of soil habitats [7] and may be able to access a wider range of carbon substrates -albeit with lower CUE [54]. Due to their metabolic diversity, bacteria with larger genomes may be more ecologically successful where resources are scarce but diverse, and where there is little penalty for slow growth [13]. In contrast, bacteria found to be resource generalists in culture have higher rRNA copy numbers, indicative of copiotroph lifestyles, than bacteria with low rRNA copy numbers that were associated with a higher degree of specialization [67]. These mixed results question whether evolutionary trade-offs between power and yield can be generalized from genomes alone. In our rhizosphere genomes, we did not find statistically significant evidence for a trade-off between resource niche breadth and CUE (Suppl. Figure S11). One possible explanation for the absence of a niche breadth-efficiency trade-off is that the energetic burden of maintaining numerous metabolic pathways is insufficiently represented by differences in genome size [33]. Alternatively, there might just not be a strong trade-off between niche breadth and growth efficiency in the rhizosphere, as environmental fluctuations in the form of the timing and composition of root exudates is a significant evolutionary driver of community succession [43, 73].

## Conclusion

Root-microbial dynamics have significant effects on the soil carbon cycle, altering the amount and types of organic matter that become associated with mineral surfaces [45]. Drawing heavily on recent studies that have identified direct predictive links between plant exudate composition and rhizosphere community assembly [56, 73], we synthesized genomic traits into model-based predictions of life history strategies for a set of rhizosphere bacteria. We found that interacting microbial traits (maximum specific growth rate, substrate uptake kinetics, ribosome biosynthesis potential and extracellular enzyme synthesis) have additional interactions with the dynamics of root exudate chemistry, creating emergent patterns of microbial carbon-use efficiency. These combinations of traits manifest as life history strategies, and have consequences for the path that small molecules take on the way to becoming stabilized soil organic matter. The trade-offs among life history strategies are not captured by traditional r-K classifications of microorganisms, or taxon-level descriptions that fail to resolve the constraining role of reserves and metabolic memory in microbial metabolism. We suggest this type of understanding is integral to more accurately representing rhizosphere dynamics in soil biogeochemical models. Our results suggest that genome-informed trait-based modeling holds great promise for connecting theoretical relationships (between genes, genomes, traits, and the environment) to biogeochemical processes that can be benchmarked rigorously with population and community scale data in future model iterations [60].

## Acknowledgements

This work was supported by the U.S. Department of Energy (DOE), Office of Biological and Environmental Research, Genomic Science Program (GSP) LLNL ‘Microbes Persist’ Soil Microbiome Scientific Focus Area SCW1632. This work was performed at Lawrence Berkeley National Laboratory funded under U.S. Department of Energy contract number DE-AC02-05CH11231. Part of this work was performed at LLNL under the auspices of the DOE, Contract DE-AC52-07NA27344.

