## Supplementary Information for "Life history strategies and niches of soil bacteria emerge from interacting thermodynamic, biophysical, and metabolic traits"

Dynamic energy budget (DEB) theory is a framework designed to model any organism [20]. The generality of DEB theory to a broad range of organism sizes, and from individuals to populations [19], stems from its scaling principles routed in classical theories of physics, such as Newton's laws of motion, Dalton's law of partial pressure, etc. This scaling coherency also makes DEB theory naturally compatible with thermodynamics [32]. The key idea of DEB models is that energy is collected from the environment and assimilated into reserve for future allocation to other components (e.g., structural biomass, maintenance, extracellular enzyme production). Maintenance, which takes priority over growth, can be paid from reserves or structural biomass depending on external resource conditions [39]. Energy allocated to growth must be converted into structure, with the yield in structure a fixed fraction of the resources invested, and with the remaining investment lost through chemical reactions needed to build the specific macromolecules that comprise the structure [21]. Both the turnover of structure and reserve biomass are density-dependent [14]. Below we present the basic equations, scaling principles and assumptions representing the DEB model as implemented in DEBmicroTrait (Suppl. Figure 1).

### A Model description

We follow the DEB convention that mass flows of compound $*$, $j_*$, are measured in moles (or C-moles) per C-mol of structural biomass per unit time [32]. Energy flows, $p_*$ are given in units of Gibbs energy per C-mol of structural biomass per unit time. Efficiencies $y_{*1*2}$ relate two mass flows and represent the number of moles of $*1$ needed to produce one mol of $*2$. Chemical potentials $\mu_*$ measured in Gibbs energy per mol or C-mol are used to convert mass to energy flows.

#### Reserve density dynamics

The change in C-mol of reserve ($M_E$) per C-mol of structural biomass ($M_V$) per unit time is determined by the difference in assimilation power ($p_A$) and mobilization power of reserve ($p_C$),

$$j_E = \frac{1}{M_V}\frac{dM_E}{dt} = \frac{p_A - p_C}{\mu_E}\,, \tag{1}$$

where $\mu_E$ is the chemical potential of reserve. Dynamics of the reserve density, $m_E = M_E/M_V$, follow from application of the chain rule of differentiation as,

$$\frac{dm_E}{dt} = \frac{p_A - p_C}{\mu_E} - m_E\frac{1}{M_V}\frac{dM_V}{dt}\,, \tag{2}$$

where $r \equiv \frac{1}{M_V}\frac{dM_V}{dt}$ is the specific growth rate. With standard DEB assumptions of (1) weak homeostasis (the reserve density is constant at constant substrate concentration and independent of structural biomass), (2) the mobilization power is independent of substrate availability, and (3) partionability of reserve, it follows that the mobilization power is given by the difference between the first order turnover rate of reserve, $k_E m_E$, and dilution by growth:

$$p_C = \mu_E(k_E m_E - m_E r)\,. \tag{3}$$

The latter term cancels in (2) and the reserve density dynamics simplify to

$$\frac{dm_E}{dt} = \frac{p_A}{\mu_E} - k_E m_E\,. \tag{4}$$

#### Specific growth rate

Reserve mobilization power is partitioned between growth, $p_G$, maintenance $p_M$, and extracellular enzyme production $p_X$, i.e.,

$$p_C = p_G + p_M + p_X\,. \tag{5}$$

The specific growth rate is then obtained by combining (5) and (3) as

$$r = \frac{p_G}{\mu_E y_{EV}} = \frac{k_E m_E - (p_M + p_X)/\mu_E}{m_E + y_{EV}} \tag{6}$$

where $y_{EV}$ is the efficiency of the growth machinery.

#### Extracellular enzyme production

We only consider free extracellular enzymes that are released into the microenvironment by the cell [40]. In the absence of regulation, extracellular enzymes are produced constitutively at a specific rate $x$ proportional to the specific growth rate, i.e.

$$x = \frac{p_X}{\mu_E y_{EX}} = \frac{z_X p_G}{\mu_E y_{EX}}\,, \tag{7}$$

where $z_X$ scales the enzyme production power according to the normalized relative frequency of glycoside hydrolase genes per genome.

#### Simulations

We simulated laboratory batch culture conditions for 39 bacterial isolates growing on 82 plant exudate metabolites. The concentration of exudates was selected to match the original growth medium concentration (125 mg C per liter), assuming that ammonium is an unlimited nitrogen source for the synthesis of biomass. Exudate concentrations correspond to concentrations of dissolved organic C detected in soil from the University of California Hopland Research and Extension Center (Hopland, CA, USA; 38°59′ 34.5768″ N, 123°4′ 3.7704″ W), where these bacteria were originally isolated. Inocula corresponding to $10^3$ cells were split into 90% reserve and 10% structural biomass [39]. Simulations were then extended to represent a mixed growth medium by evenly distributing the original batch exudate concentration across the different metabolites.

### B Membrane requirements for substrate uptake at balanced growth

In analogy to enzyme kinetics, the equilibrium chemistry approximation (ECA) for substrate uptake [35] assumes that the maximum specific uptake rate, $V_{max}$, scales linearly with substrate binding-site density on the cell surface, whereas the saturation of uptake capability with increasing binding-site density is reflected in half-saturation constants $K$ [37]. We estimated membrane requirements for substrate uptake in DEBmicroTrait under conditions of balanced growth. Then, the ratios of extensive properties (i.e., intensive properties, such as the reserve density) remain constant [4]. From Eq. 4 it follows that the reserve density is given by the ratio of assimilation power to reserve turnover rate

$$m_E^* = \frac{p_A^*}{\mu_E k_E}, \tag{8}$$

where the superscript * refers to balanced growth conditions. Here, theory provides an extra degree of freedom since maximum assimilation power, reserve turnover rate, and reserve density all seem appropriate to take as primary parameters that may be determined independently through natural selection. While it has become standard in the DEB literature to take $m_E^*$ and $k_E$ as the primary parameters, this assumption ultimately leads to larger organisms with similar $k_E$ having larger maximum reserve densities, which seems counterintuitive [21]. At balanced growth, cells have evolved optimal protein densities in cellular compartments, e.g. the cell membrane, that maximize reaction rates [8, 18]. Hence, a common assumption in the modeling literature is that the cellular membrane area that needs to be covered with binding sites is optimized to maximum specific growth rates [12, 26]. Solving Eq. 6 and Eq. 8 for the substrate-specific binding site density in the ECA framework [37] shows that the fraction of the membrane area required for near-maximal metabolite assimilation rates ranges from 0.0005% to 0.19%, with a median of 0.1%. The average estimated binding-site density for cumulative uptake of plant metabolites is $\approx 0.08$, or 8% of the total membrane area [2, 26].

In order to test our estimates, we independently benchmarked predictions of half-saturation constants using a compilation of existing data that explicitly differentiates between transport and growth kinetics of bacteria [5]. Our predictions were based on reference genomes and median rrn operon copy number at genus level under the NCBI taxonomy [33]. In the absence of reported maximum specific growth rates, the maximum specific transport rates were used to estimate a commensurate binding-site density. The predicted half-saturation constants did not differ significantly from the majority of measured values (Suppl. Figure 1). Upon excluding two measurements that were performed via flow dialysis instead of radioactivity uptake with incubation times less than ten minutes, the model $r^2$ was 0.82. By further removing the value with largest residual, corresponding to *Bradyrhizobium japonicum* with very large reference genome size (9.32 Mbp) and low rrn copy number (2), the model $r^2$ improved to 0.92, with intercept (p=0.47) and slope (p=0.42) that did not differ significantly from the 1-1 line. The unexplained variance is likely due to strain level variation in genomic trait parameters that could not be accounted for in the NCBI database.

In addition to $V_{max}$ and $K$, the specific substrate affinity ($V_{max}$ divided by $K$) is a good measure for comparing interspecies competitiveness at low substrate concentrations [6]. While pure diffusion limitation suggests that affinities scale linearly with cell radius, thus conferring fitness advantages to biophysically efficient cell morphologies with large effective linear dimensions [42], higher density requirements for small cells, together with higher relative investment costs for binding proteins, might explain why observed affinities for single nutrient uptake scale approximately with the square of the cell radius [15, 22], i.e., cell surface area (Suppl. Figure 2 B). Evolutionary history, however, can obscure the biophysical advantage of small cells at low nutrient concentrations: upon allocating binding sites according to relative gene frequencies of transporter genes (Suppl. Table 1), surface area-to-volume ratios and genomic investment explain 38%, respectively 14%, of the variance in specific affinities across different metabolites and isolates.

### C Ribosome requirements and protein overexpression

The growth rate of bacteria is strongly tied to the relative abundance of macromolecular components, including ribosomes, RNA and proteins [4]. Previous efforts have characterized the scaling of major macromolecular components across a wide range of bacterial cell sizes [16]. According to [16], the cytoplasmic cell volume taken up by expressed proteins, $V_p$, scales sublinearly with total cellular volume, $V_c$, following

$$V_p = P_0 V_c^{\beta_p}, \tag{9}$$

where $P_0$ = 3.42e-7 (m$^3$ protein (m$^3$ cell)$^{-\beta_p}$, and $\beta_p = 0.70 \pm 0.06$. The total protein content constrains the number of ribosomes, $N_r$, and the overall RNA content of the cell, according to the following inequality,

$$N_r \geq \frac{\bar{l}_p N_p(\phi/r + 1)}{\bar{r}_r/r - \bar{l}_r(\eta/r + 1)}, \tag{10}$$

where r is growth rate, $\bar{l}_r$ is is the average length of a ribosome in base pairs, $\bar{r}_r$ is the maximum base pair processing rate of the ribosome, $\bar{l}_p$ is is the average protein length found to be invariant across bacteria [45], $\eta$ and $\phi$ are specific degradation rates for ribosomes and proteins, and $N_p$ is the total number of proteins. Protein and ribosome numbers were converted to volumes by multiplying with the volume, $\bar{v}_p$ and $\bar{v}_r$, of an average protein or ribosome, respectively. Following [13], we conceptualized the specific reserve turnover rate $k_E$ as the maximum synthetic capacity of the growth machinery, synonymous with 'translation power', i.e. the rate of protein synthesis in a cell or culture at maximum specific growth rate normalized to the biomass invested in the protein synthesis system [9]. Thus, $k_E$ is given by

$$k_E = \frac{r_{max} P}{R}\,, \tag{11}$$

with maximum specific growth rate $r_{max}$, protein mass $P$, and rRNA mass $R$ as a proxy for the biomass invested in the protein synthesis system. In order to evaluate the allometric model for $k_E$, we regressed observed values of translational performance in culture [9] against predicted values and compared slope and intercept parameters against the 1:1 line ( [27], Suppl. Figure 3 A). Intercept ($p = 0.25$) and slope ($p = 0.07$) did not differ significantly from 0 and 1. Hence most of the model prediction errors were due to unexplained variance ($r^2 = 0.78$). The root mean squared deviation of 0.75 1/h was less than 7% of the total variation in protein translation power across the range of observed cell sizes. Translation power is significantly higher in isolates with a positive response to root growth (p=0.02). Based on existing protein synthesis phenotypes [28], we used the scaling relationship with ribosomal RNA operon copy number to estimate the corresponding translation efficiency of isolates, showing that positive responders are trending towards higher efficiency as compared to negative responders (p=0.06, Suppl. Figure 3 B). Previous modeling efforts show that protein overexpression can significantly lower biomass yield [25, 44]. Similarly, we find that the maximum structural biomass yield, defined as the fraction of reserve that is mobilized for growth, i.e.,

$$y_V = \frac{r}{k_E m_E} \tag{12}$$

is negatively correlated with $k_E$ at high $k_E$ (Suppl. Figure 3 C). As a result, maximum growth rates can occur at sub-optimal yield. At low $k_E$, the structural biomass yield remains relatively constant because reserves can be mobilized efficiently (Suppl. Figure 3 D).

### D Mass and energy coupling in DEB metabolism

In DEBmicroTrait, the substrate assimilation into reserves and the subsequent transformation of reserves into structural biomass are subject to stoichiometric constraints [19]. Both substrates and reserve have a dual role: they are used to drive the transformation and they serve as building blocks (anabolic substrates). The thermodynamic electron equivalents model (TEEM) considers the energy provision of catabolic reaction ($\Delta G_{cat}$) to meet the energy spent in anabolism that likewise consists of (1) the conversion of carbon source to biomass building blocks ($\Delta G_{block}$), and (2) the conversion of biomass building blocks to biomass ($\Delta G_{syn}$) [24]. The energy balance can be written as

$$-\lambda\left(\eta \Delta G_{cat}\right) = \eta^m \Delta G_{block} + \Delta G_{syn}, \tag{13}$$

where $\eta$ is a fixed energy transfer efficiency parameter for all enzymatic steps, and $m = \pm 1$ depending on whether energy is generated ($\Delta G_{block} < 0$) or consumed ($\Delta G_{block} > 0$) in the process of converting the carbon source to biomass building blocks. The parameter $\lambda$ implies how many times the catabolic reaction needs to run in order to produce sufficient Gibbs energy for the synthesis of a unit C-mole of biomass. Following [31], we assumed that the composition of biomass and building blocks is constant and equal across isolates (the strong homeostasis assumption [32]). With the value of $\lambda$ calculated as such, one can then solve the full stoichiometric equation by coupling the catabolic and anabolic reaction in DEB metabolism,

$$y = \lambda y^{cat} + y^{block}\,, \tag{14}$$

requiring only the identification of the anabolic carbon and nitrogen source, and the electron donor-acceptor couple in the energy generating catabolic reaction [17]. The substrate assimilation yields across different metanolite classes are shown in Suppl. Figure 4.

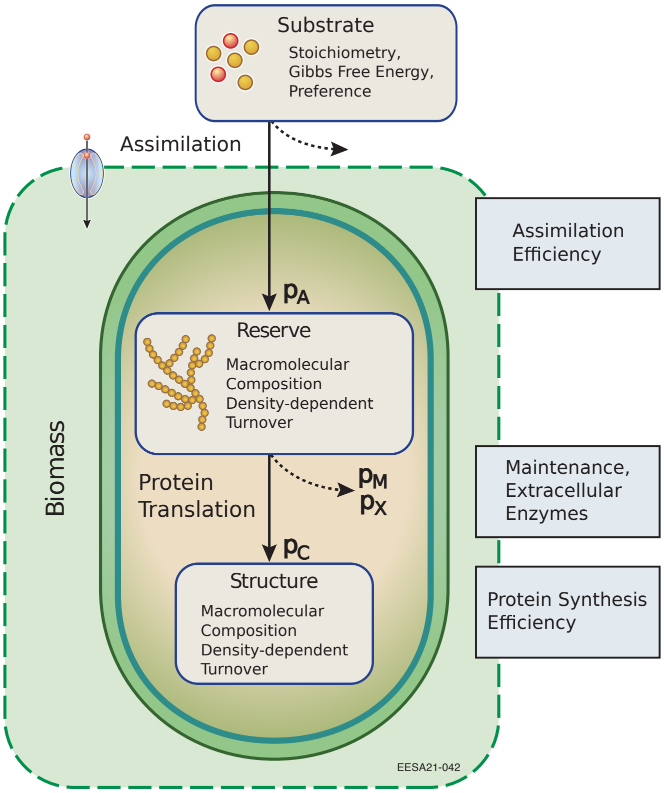


**Figure 1.** Dynamic energy budget allocation for a single-reserve, single-structure heterotrophic microorganism feeding on a single substrate. Substrate assimilation occurs via uptake through membrane transporters. Dotted arrows represent energy flows diverted from growth, either due to assimilation and protein synthesis inefficiencies, maintenance or the production of extracellular enzymes.

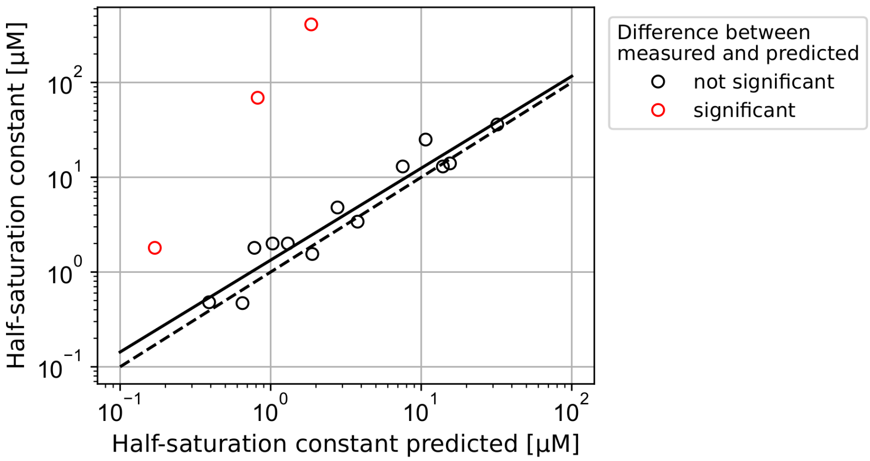


**Figure 2.** Comparison of predicted half-saturation constants and reported half-saturation constants [6] for 11 reference genomes. The solid line indicates the regression line ($r^{0.92}$). The dashed line corresponds to the 1-1 line for observed vs. predicted values. Predictions with significant differences from the 1-1 line are denoted by red color.

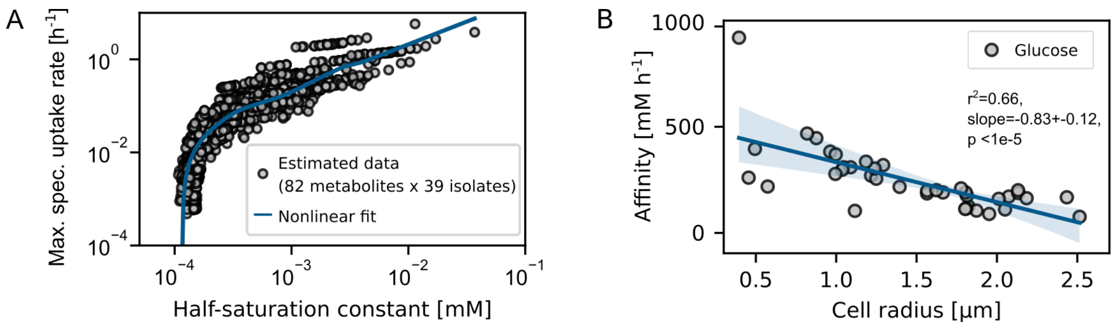


**Figure 3.** Trade-offs and scaling laws in substrate uptake of isolates. **A** Trade-off curve between maximum specific uptake rate and half-saturation constant across metabolites (n=82) and isolates (n=39). **B** Power law relation between membrane binding site density and isolate cell radius for glucose uptake. The blue line indicates the observed regression line, while the shaded area indicates the 95% confidence band. The slope with standard error, p-value, and the corresponding $r^2$ of the regressions are shown.

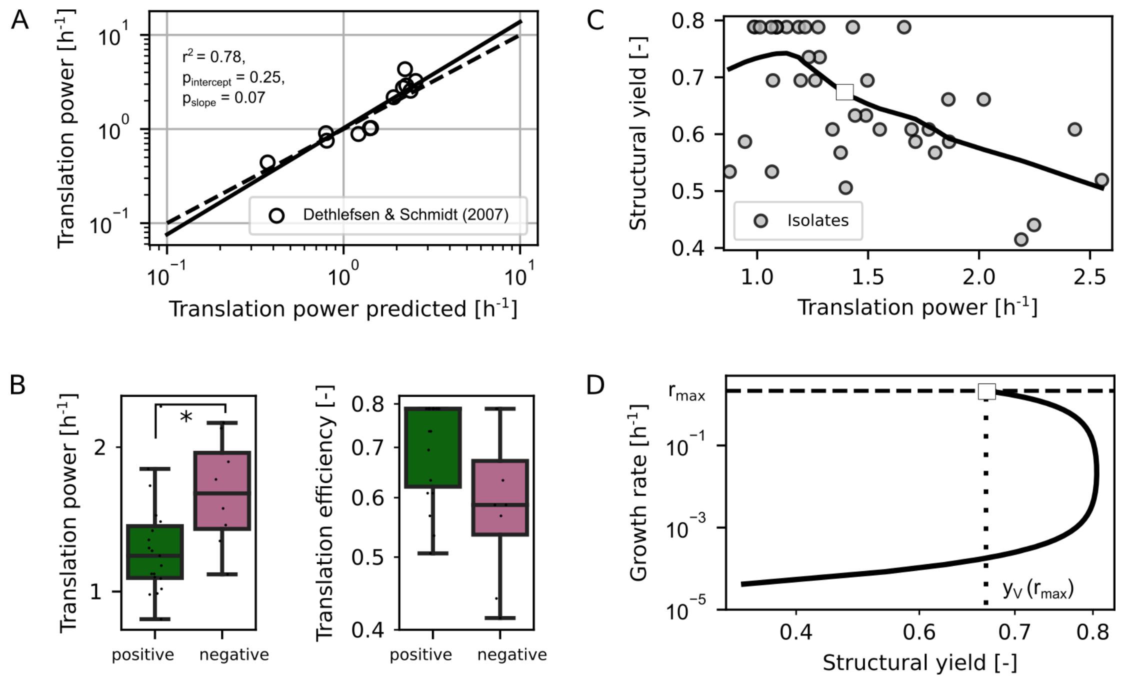


**Figure 4.** Protein synthesis phenotypes and implications for rate-yield trade-offs. **A** Regression scatter plot for protein synthesis power. The dashed line indicates the regression line. The solid line corresponds to the 1-1 line for observed vs. predicted values (n=12). **B** Protein synthesis power and efficiency of soil isolates. Differences in trait distributions between positive and negative responders were evaluated using the Kruskal-Wallis one-way analysis of variance. In each boxplot, a point denotes a single isolate. The top and bottom of each box represent the 25th and 75th percentiles, the horizontal line inside each box represents the median and the whiskers represent the range of the points excluding outliers. **C** Relationship between structural biomass yield and translation power across isolates (n=39). The blue line corresponds to a locally weighted linear regression model. **D** Exemplary relationship between growth rate and yield (n=1). The dotted line marks the sub-optimal structural yield value at maximum specific growth rate (dashed line).

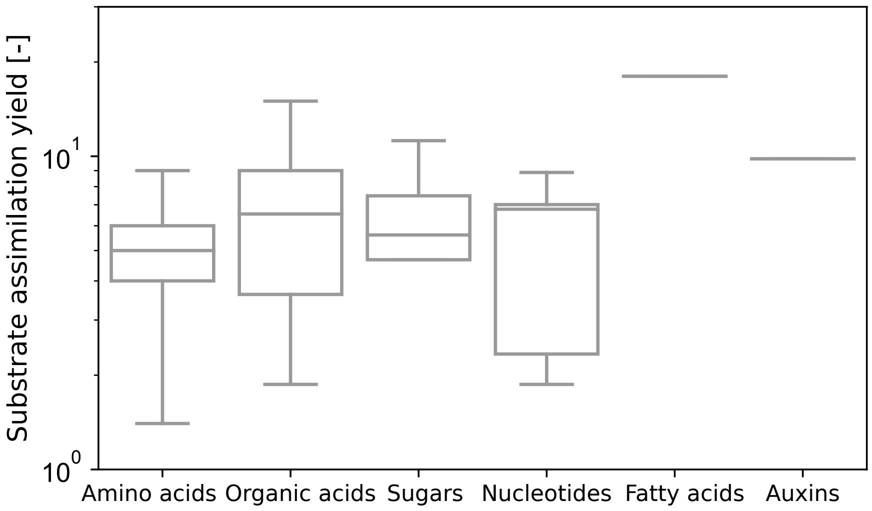


**Figure 5.** Substrate assimilation yield of isolates across root metabolite class. The top and bottom of each box represent the 25th and 75th percentiles, the horizontal line inside each box represents the median and the whiskers represent the range of the points excluding outliers.

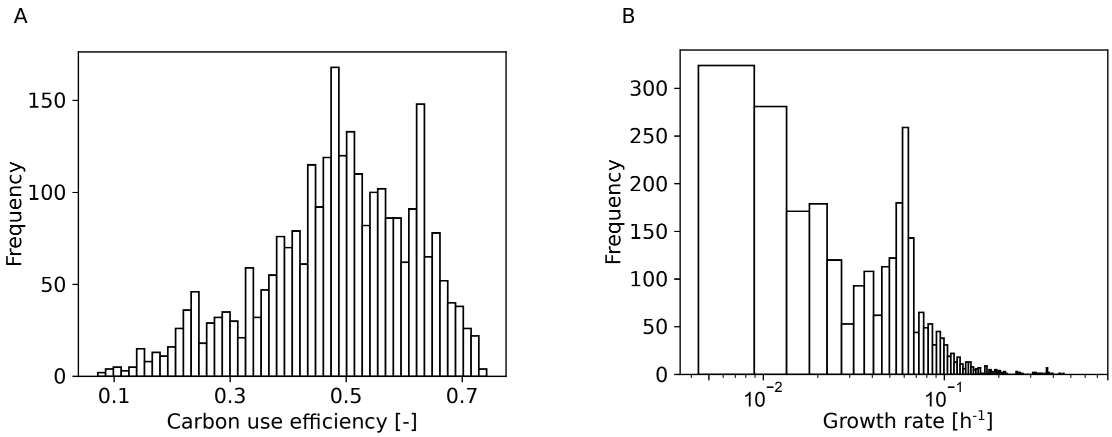


**Figure 6.** Histogram of (A) predicted carbon use efficiency and (B) predicted growth rate across isolates (n=39) and plant metabolites (n=82) in DEBmicroTrait batch simulation.

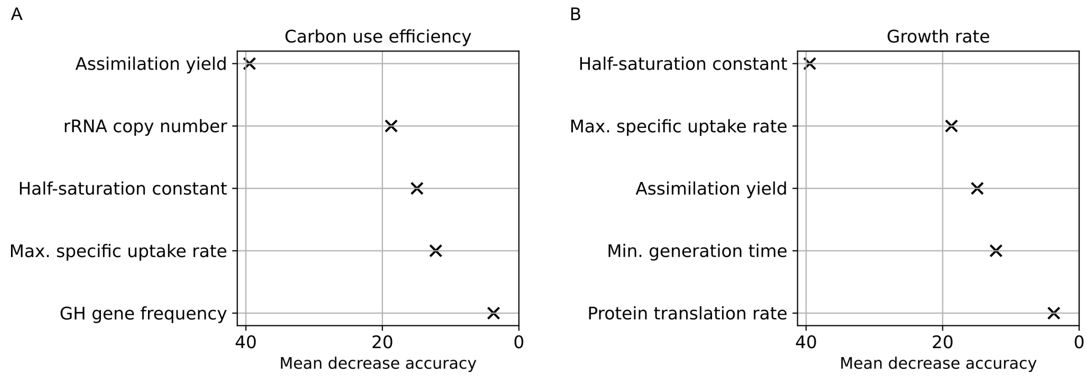


**Figure 7.** A representation of the five most influential predictors of carbon-use efficiency (A) and growth rate (B) in batch simulations as determined by mean decrease in accuracy. The mean decrease in accuracy is a measurement of the change in the accuracy of the random forest's predictions when the variable in question is randomly permuted. Labels on the y axis indicate the feature names. Feature contributions for all case studies were computed on predictions for the undefined rhizosphere response group as out-of-bag samples using the forestFloor package available in R [43].

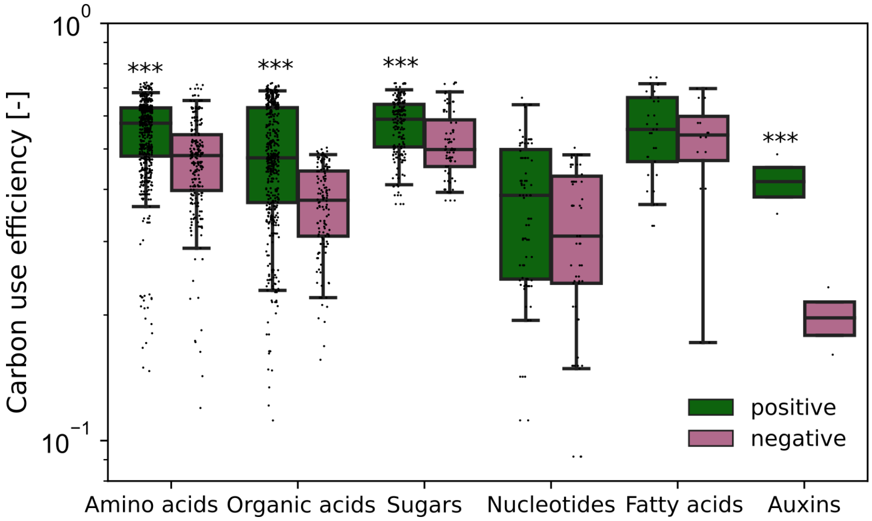


**Figure 8.** Predicted carbon-use efficiency of isolates. Differences in distributions between positive and negative responders were evaluated using the Kruskal-Wallis one-way analysis of variance. The top and bottom of each box represent the 25th and 75th percentiles, the horizontal line inside each box represents the median and the whiskers represent the range of the points excluding outliers. Substrates: n=82, consumers: n=27.

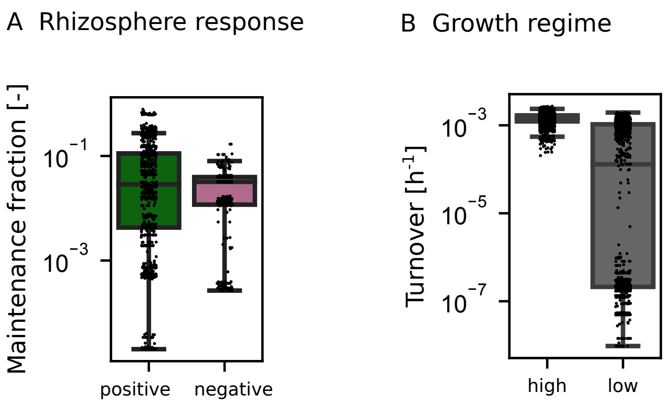


**Figure 9.** Selection for efficiency at low growth rates. (A) The fraction of the total energy budget invested in maintenance depending on rhizosphere response (positive, negative) in the low growth rate regime (p=2.1e-5). (B) Emergent biomass turnover in the high and low growth rate regime. The distribution in the low growth rate regime is bimodal, with some organisms with significantly lower turnover rates (p<1e-12). Differences in distributions between positive and negative responders were evaluated using the Kruskal-Wallis one-way analysis of variance. The top and bottom of each box represent the 25th and 75th percentiles, the horizontal line inside each box represents the median and the whiskers represent the range of the points excluding outliers.

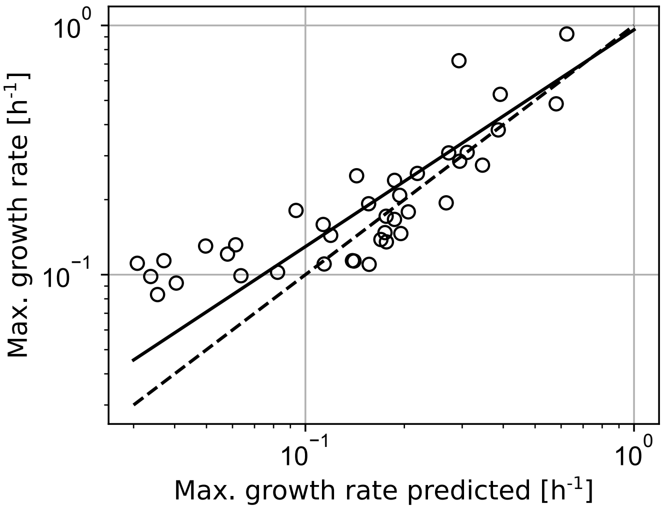


**Figure 10.** Comparison of predicted maximum specific growth rates of Hopland isolates (n=39) and confirmed genome-predicted maximum specific growth rates through laboratory growth rate experiments [46]. The solid line indicates the regression line ($r^2$=0.67). The dashed line corresponds to the 1-1 line for observed vs. predicted values.

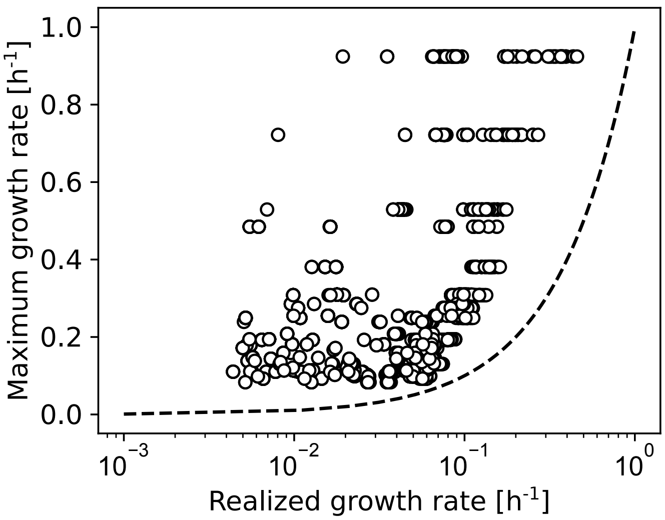


**Figure 11.** Comparison of realized growth rates for growth on root exudate metabolites (n=82) and genome-predicted maximum specific growth rates of Hopland isolates (n=39). The x-axis was log scaled. The dashed line corresponds to the 1-1 line.

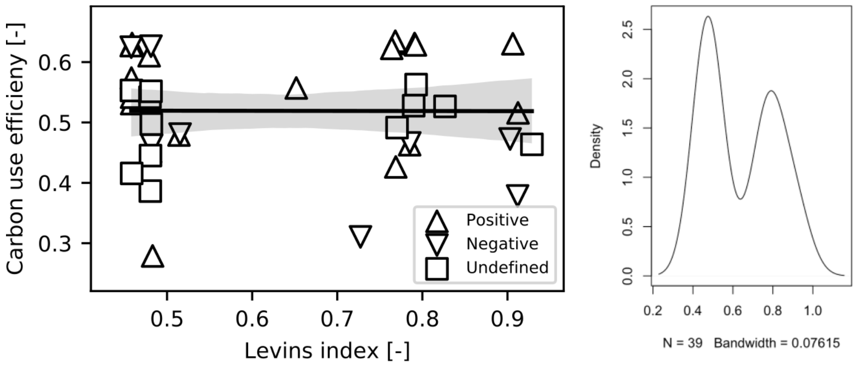


**Figure 12.** Relationship between resource niche breadth (Levins index, [7]) and carbon-use efficiency of isolates across root metabolites ($F_{1,37} < 3e^{-4}$, $r^2 < 1e^{-5}$, $p = 0.99$). Niche breadth was unimodally distributed across isolates based on Hartigan's dip test ($D = 0.12$, $p < 2e^{-5}$), both in the low ($D = 0.13$, $p < 7e^{-4}$) and high ($D = 0.15$, $p < 2e^{-3}$) growth regime (not shown).

**Table 1.** Taxonomic and substrate variance partitioning using linear mixed-effects models describing the effect of isolate identity, taxonomic order, metabolite type and metabolite class on carbon use efficiency.

| Model | Main variable | Nested variable | Variance explained (main) | Variance explained (nested) | AIC |
|---|---|---|---|---|---|
| **Taxonomic order** | Species | Metabolite | 38% | 88% | -8624 |
| | Phylum | | 13% | 63% | -5754 |
| | Class | | 20% | 69% | -6222 |
| **Metabolite type** | Metabolite | Species | 48% | 88% | -8553 |
| | Class | | 15% | 54% | -5465 |

Mixed-effects models were fit by REML using the lmer() function in the lme4 R package. In these models there is a main variable and a nested variable. For each analysis, the variation explained by the main variable is accounted for before the variation explained by the nested variable is determined. As such, these results indicate the relative importance of each variable when grouped together in a nested framework.

**Table 2.** Model selection for predicting isolate growth rates and carbon use efficiency based on rRNA copy number (rrn) and genome size (G). The predictions were split based on growth rate into a high (>0.041 $h^{-1}$) vs. low (<0.04 $h^{-1}$) growth regime.

| Growth regime | Model | Slope rrn | Slope G | Intercept | p-value rrn | p-value G | p-value rrn:G | $r^2$ | AIC |
|---|---|---|---|---|---|---|---|---|---|
| **Growth rate** | | | | | | | | | |
| High | rrn | 0.0125 | | 0.053 | <2e-16 | | <2e-16 | 0.30 | **-4997** |
| | G | | -1.764e-8 | 0.165 | | <2e-16 | <2e-16 | 0.23 | -4847 |
| | rrn:G | 0.0131 | 1.148e-09 | 0.0459 | <2e-16 | 0.503 | <2e-16 | 0.30 | -4996 |
| Low | rrn | 0.00117 | | 0.0168 | 4.53e-8 | | 1.17e-10 | 0.04 | **-8427** |
| | G | | -8.335e-10 | 0.0228 | | 0.00015 | 0.13467 | 0.02 | -8411 |
| | rrn:G | 0.00124 | 1.027e-10 | 0.0162 | 7.7e-5 | 0.749 | 2.90e-05 | 0.04 | -8425 |
| **Carbon-use efficiency** | | | | | | | | | |
| High | rrn | -0.0148 | | 0.628 | <2e-16 | | 0.641 | 0.44 | **-3089** |
| | G | | 2.216e-08 | 0.488 | | <2e-16 | <2e-16 | 0.42 | -3051 |
| | rrn:G | -0.0128 | 3.745e-09 | 0.605 | 3.66e-10 | 0.263 | 0.44 | 0.44 | -3088 |
| Low | rrn | -0.0184 | | 0.582 | 3.99e-14 | | 0.807 | 0.35 | -1952 |
| | G | | 9.641e-9 | 0.505 | | 0.000127 | < 2e-16 | 0.32 | -1909 |
| | rrn:G | -0.0249 | -9.170e-9 | 0.636 | 2.81e-12 | 0.0117 | 0.0634 | 0.35 | **-1956** |

All models have the generic form: Dependent variable = copy number * slope rrn+genome size * slope G + copy number*genome size *slope rrn:G+intercept. Blank cells indicate cases where a term was excluded from the model (e.g., a model based on rrn, rRNA copy number, will not have a slope or p-value estimate for G, genome size). For cases where at least one model was statistically significant, the best model based on the smallest AIC value is indicated in bold.

**Table 3.** Genomic traits of bacterial isolates used to constrain DEBmicroTrait.

| Parameter | Symbol/Formula | Description \| Unit | Reference |
|---|---|---|---|
| **Genomic traits** | | Genome sequences of isolates were analyzed for the specific genomic traits listed below as described in more detail in the listed reference. | [46] |
| Minimum generation time | $\frac{\mathrm{ENC_{all}}-\mathrm{ENC_{ribosomal\ protein\ genes}}}{\mathrm{ENC_{all}}}$ | Predicted based on codon-usage bias between all genes and a set of highly expressed (ribosomal protein) genes following a linear regression model (ENC; effective number of codons given G + C composition) \| [$h^{-1}$] | [41] |
| Genome size | $L_{DNA}$ | \| [bp] | |
| 16S rRNA copy number | | \| [-] | |
| Glycoside hydrolase genes | $z_X$ | Normalized gene copy numbers of glycoside hydrolase genes were predicted using a hidden Markov model search against the Carbohydrate Active enZYmes database (CAZy). \| [-] | [10] |
| Transporter genes | $z_B$ | Normalized gene copy numbers of transporter genes were predicted using a hidden Markov model search against the TransportDB database. \| [-] | [11] |

**Table 4.** Ontogenic growth model parameters and scaling relationship used to constrain DEBmicroTrait.

| Parameter | Symbol/Formula/Value | Description \| Unit | Reference |
|---|---|---|---|
| **Macromolecular composition** | | | [16] |
| $V_{DNA}$ | $V_{DNA} = v_N L_{DNA}$ | Genome volume \| [m$^3$] | |
| $v_N$ | 1.47e-27 | Nucleotide volume \| [m$^3$] | |
| $V_c$ | $V_c = (V_{DNA}/D_0)^{1/\beta_D}$ | Cell volume \| [m$^3$] | |
| | $D_0$ = 3e-17, $\beta_D$ = 0.21 | | |
| $V_p$ | $P_0 V_c^{\beta_p}$ | Protein volume \| [m$^3$] | |
| | $P_0$ = 3.42e-7, $\beta_P$ = 0.70 | | |
| $v_p$ | 4.24e-26 | Protein volume \| [m$^3$] | |
| $N_r$ | $\frac{\bar{l}_p N_p(\phi/r+1)}{\bar{r}_r/r-\bar{l}_r(\eta/r+1)}$ | Ribosome number \| [-] | |
| $\bar{l}_p$ | 975 | Average protein length \| [bp] | |
| $N_p$ | | Number of proteins \| [-] | |
| $r$ | | Population growth rate \| [s$^{-1}$] | |
| $\phi$ | 6.2e-5 | Specific protein degradation rate \| [s$^{-1}$] | |
| $\eta$ | 6.2e-5 | Specific ribosome degradation rate \| [s$^{-1}$] | |
| $\bar{l}_r$ | 4566 | Average ribosome length \| [bp] | |
| $\bar{r}_r$ | 63 | Maximum ribosome processing rate \| [bp s$^{-1}$] | |

**Table 5.** DEBmicroTrait model parameters

| Parameter | Symbol/Formula/Value | Description \| Unit | Reference |
|---|---|---|---|
| **Assimilation** | | | |
| $\rho_b$ | | Membrane binding site density \| [-] | |
| $m_C$ | 12.011 | Molecular weight of carbon \| [g mol$^{-1}$] | |
| $r_b$ | 1e-9 | Binding site radius \| [m] | [34] |
| $k_{2,p}$ | 180.0 | Maximum substrate conversion rate \| [s$^{-1}$] | [26] |
| $m_d$ | $\sum_i m_{comp,i}$ | Cellular dry mass \| [g] | |
| $m_{comp}$ | | Macromolecular component mass \| [g] | [16] |
| $n_0$ | $\frac{m_C}{0.47 m_d N_A}$ | Cell number density (assuming 47% carbon per dry mass) \| [mol cell gC$^{-1}$] | [30] |
| $N_{SB}$ | $\frac{n_0 4\pi r_c^2 \rho_b}{m_C \pi r_b}$ | Binding site density \| [mol sites (mol biomass C)$^{-1}$] | [37] |
| $p_{int}$ | $\frac{4\rho_b r_c^2}{r_b(4\rho_b r_c^2/r_p+\pi r_c)}$ | Substrate interception probability | [2] |
| $K$ | $\frac{k_{2,p}\rho_b r_c}{D_S \pi r_b^2 N_A} p_{int}^{-1}$ | Binding half-saturation constant | [38] |
| $N_C$ | | Number of carbon atoms \| [-] | |
| $y_{DE}$ | $\frac{1}{y^{an}+\lambda y^{cat}}$ | Assimilation yield | [17] |
| $y^{an}$ | | Stoichiometric vector of anabolism | |
| $y^{cat}$ | | Stoichiometric vector of catabolism | |
| $\lambda$ | | Coupling parameter between catabolism and anabolism | |
| **Core metabolism** | | | |
| $k_M$ | $0.39 V_c.^{0.88}$ | Basal maintenance rate \| [h$^{-1}$] | [23] |
| $y_{EM}$ | 1.0 | Fraction of the maintenance costs paid from reserve | [39] |
| $k_E$ | $\frac{r_{max} V_p}{V_r}$ | Translation power | [9] |
| $y_{EV}$ | $9.5 - 1.22 \log_2(rrn)$ | Translation efficiency | [29] |
| $\alpha_X$ | 1e-2$z_X$ | Constitutive enzyme expression | [40] |
| **Turnover** | | | |
| $\gamma_{V,0}$ | $0.23 e^{0.88 r_{max}}$ | Max. turnover rate \| [h$^{-1}$] | [3] |
| $\gamma_{V,1}$ | $N_{cells}$*1e6*$\rho_{bulk} * m_d/m_C$ | Turnover half-saturation constant \| [$\mu$M] | [36] |
| $\rho_{bulk}$ | 1.6 | Soil bulk density \| [g ml$^{-1}$] | [1] |
